# Unveiling the Holliday junction conformational dynamics of upon binding with DNA architectural proteins IHF; A Single molecule FRET approach

**DOI:** 10.1101/2023.09.25.559264

**Authors:** Farhana Islam, Padmaja P. Mishra

## Abstract

Integration host factor (IHF) of *E. coli* is a nucleoid-associated protein with diverse roles in DNA packaging, viral DNA integration, and recombination. IHF binds to duplex DNA containing a 13 bp consensus sequence with nanomolar affinity and induces a significant bend of approximately 160° upon binding. While the Wild type IHF (WtIHF) is involved in DNA bending, thereby facilitating the integration of foreign DNA into the host genome, its engineered version, Single chain IHF (ScIHF) was designed for specific genetic engineering and biotechnological applications. We investigated interaction of the two IHF variants with Holliday junctions (HJ), crucial intermediates in DNA repair and homologous recombination. Our finding demonstrate that both variant of IHF binds to HJs with high affinity in presence of the consensus sequence, indicating a structure-based recognition mechanism. HJs are dynamic structures that can adopt open or stacked conformation. The open conformation facilitates processes like branch migration and strand exchange. Through quantitative binding studies using microscale thermophoresis, we determined the binding of IHF to four-way DNA junctions that harboured two specific binding sequences H’ & H1. Circular dichroism (CD) experiments revealed the protein’s impact on the junction conformation. This was further confirmed by Single molecule Förster resonance energy transfer (smFRET) technique that was used to examine the binding of IHF to the junction and its effect on the dynamicity of junction conformation. We also probed the population distribution of junction conformations. Interestingly, our results revealed that binding of both WtIHF & ScIHF shifts the population towards the open conformation of the junction and stabilised it in that conformation. In summary, our findings demonstrate that IHF binds HJs with a strong affinity and has a stabilizing effect on maintaining the junction’s open conformation.

## Synopsis

Holliday junctions are dynamic DNA secondary structures that are crucial for maintenance of genetic stability and facilitating genetic diversity. The widespread occurrence of Holliday junction in important cellular processes makes them suitable target for several regulatory and architectural and histone-like proteins. Integration host factor is one such abundant nucleoid associated protein found in bacterial cells that take part in site specific recombination. With the help of ensemble techniques and SmFRET, we have probed the interaction between IHF and HJ and subsequently the dynamics of HJ upon IHF binding.

- HJs adopt different conformations under different ionic conditions. CD and SmFRET show that while the open state is favoured at low concentrations of monovalent ion, upon addition of divalent cation like magnesium, the stacked form is favoured.
- For both Mg^2+^ concentrations tested (2mM and 50mM), the two junction constructs predominantly favoured the isoⅠ conformer.
- Both WtIHF & ScIHF bind to HJs with high affinity even in presence of 50mM MgCl_2_.
- Single-molecule FRET of HJ reveals conformational dynamics associated with IHF binding.
- WtIHF and ScIHF upon binding to HJs impart conformational stability to junctions in the open state even in presence of magnesium.

## Introduction

Integration Host Factor (IHF) emerges as a pivotal participant within the intricate synchronization of cellular genetics, coordinating a multifaceted ensemble of processes encompassing transcription, recombination, replication, and viral integration. Initially discovered for its role in λ-DNA integration (Nash & Robertson, 1981), IHF’s influence extends to the Cas1-Cas2 integrase system (Nuñ Ez *et al*, 2016), where it manifests as a catalytic enhancer, heightening integration precision and efficiency through pronounced perturbation of the leader sequence. This protein exhibits a strong affinity for DNA duplexes that possess a 13 bp consensus sequence, WATCARNNNNTTR (W represents A or T, R represents A or G, and N denotes any nucleotide). Upon binding, IHF invokes a conspicuous torsional aberration with in the DNA structure (Lorenz *et al*, 1999), An X-ray co-crystal structure of IHF-DNA complex reveals a bend angle approximating 160° (Rice *et al*, 1996). The recognition of IHF’s consensus sequence entails a fusion of direct and indirect readout mechanisms. IHF and the structurally kin HU being the constituents of the DNABII protein family, acknowledged as architectural proteins due to their ability to bind with high affinity to distorted DNA substrates, thereby effectuating DNA bending. Intriguingly, their capabilities extend to DNA compaction, potentially influencing gene expression via modulation of supercoiling and genome architecture (Dorman, 2015, 2013). Despite structural homology, HU and IHF diverge in DNA binding properties, the former serves as a non-sequence-specific DNA binding protein (Amemiya *et al*, 2021). Both proteins play crucial roles in bacterial biofilms, where extracellular DNA (eDNA) underpins the matrix component (Devaraj *et al*, 2015, 2018). The branched eDNA matrix harbours Holliday Junction-like structures that are bound and stabilized by cooperative influence of IHF and HU (Devaraj *et al*, 2019). Whileprevious investigations have unveiled HU’s interactions with Holliday Junctions (Bonnefoy *et al*, 1994), direct evidence substantiating IHF’s binding with these structures remains an ongoing pursuit.

Holliday Junctions (HJ) play essential roles as intermediates in DNA repair and recombination processes (Liu & West, 2004; Spies & Fishel, 2015) and have recently been identified as structural components of pathogenic biofilms (Devaraj *et al*, 2019). These dynamic structural motifs exhibit a propensity of structural flexibility, existing in diverse conformations (Lilley, 2000; Song *et al*, 2022). The structural details of HJs, as revealed by X-ray crystallography studies, manifests as an intersection of four DNA strands and exchanging in a central region (Eichman *et al*, 2000). This central region, characterized by a high negative charge, has been observed to accommodate ions structurally (Møllegaard *et al*, 1994; Murchie *et al*, 1989). The open conformation facilitates branch migration, while the stacked counterpart surviving as a prime target for enzymatic intervention by resolvases and endonucleases (Mahmoud & Dhakal, 2022; Lilley, 2017; Wyatt & West, 2014). Junction conformation plays an important role in branch migration as it has been found that the open extended form of the junction, that is found under low salt conditions, facilitate branch migration (Panyutin *et al*, 1995; Karymov *et al*, 2005). Notably, the junction often oscillates between an open and a stacked-X conformer (Mckinney *et al*, 2002; Grainger *et al*, 1998; Promisel *et al*, 1989), the former spreading the arms wide apart to minimize electrostatic repulsion along the DNA backbone. However, presence of divalent cations engenders an anti-parallel stacked-X conformation, symptomatic of electrostatic charge neutralization by metal ions (Duckett *et al*, 1990). The presence of magnesium ions, acting as a regulatory conductor, results in a decrease in branch migration rate and depopulation of open state. Earlier studies deploying single-molecule FRET have unravelled intricate state transitions and conformational heterogeneity intrinsic to HJ dynamic, highlighting the interplay between internal structural diversity and the modulating effect of Mg^+2^ ions (Joo *et al*, 2004). Notably, enzymes participating in branch migration or nucleolytic resolution tend to stabilize the planar symmetrical structure as physiologically relevant (Lilley, 2017; Sarbajna & West, 2014; Ray *et al*, 2021; Sobhy & Abdelmaboud). Foremost among techniques unravelling IHF-DNA interactions is X-ray crystallography, a pivotal medium capturing the three-dimensional structure of IHF in complex with DNA at atomic resolution (Yang & Nash, 1989). Multiple crystal structures have indicated the nature of IHF-DNA interaction, accompanied by conformational perturbation on complexation (Bao *et al*, 2004; Lynch *et al*, 2003). IHF is found to bind to DNA as a homodimer, with each monomer consisting of two structurally similar domains connected by a flexible linker. The N-terminal domains are responsible for DNA recognition and binding, while the C-terminal counterpart engages in protein-protein interactions and DNA bending. The binding of leads to a prominent twist in the DNA helix, facilitating the protein-protein ensembles and promoting DNA looping. NMR spectroscopy has provided insights into the dynamic behaviour of IHF and its interactions with DNA in solution (Zhou *et al*, 2019; Dhavan *et al*, 1999). Electron microscopy studies have visualized IHF-DNA complexes at lower resolutions, revealing the overall architecture of the complexes and the global DNA bending induced by IHF. Single-molecule techniques, including atomic force microscopy and optical tweezers, shed light on the mechanical properties of IHF-DNA complexes (Bera *et al*, 2018). These studies showcase the force-dependent binding and DNA bending dynamics, shedding light on the energetics and kinetics of these processes. These endeavours uncover IHF’s multivalent role in DNA recombination and gene regulation., establishing its status as a critical participant (Goosen & van de Putte, 1995; Boccard & Prentki, 1993; Reverchon *et al*, 2021). In recombination, IHF serves as a cofactor to recombinases, facilitating the integration or excision of DNA segments. Moreover, IHF’s involvement in gene expression modulation through targeted binding to specific regulatory regions in the bacterial genome is evident, thereby modulating the accessibility of the DNA to transcription factors and RNA polymerase. The results from the biophysical studies on IHF and DNA augments our comprehension of both structural and functional facets. These studies have advanced our understanding of the mechanisms underlying DNA bending, protein-DNA recognition, and gene regulation, with potential implications for the development of novel therapeutic strategies targeting bacterial pathogens. Considerable progress has been achieved in elucidating the structure and functional attributes of integration host factor (IHF) and its interactions with DNA. However, several knowledge gaps persists. While IHF’s capability to induce substantial bending to DNA upon binding is acknowledged, the precise mechanism the precise mechanism governing the phenomenon remain incompletely understood. Further studies are imperative to unravel the molecular basis of IHF-induced DNA bending, necessitating identification of the specific amino acid residues and underlying structural features involved in this process. Importantly, most biophysical studies conducted on IHF and DNA have focused on static structures or averaged measurements, there by offering limited insight into the dynamics and conformational flexibility of IHF-DNA complexes. A holistic understanding demands exploration of the dynamic behaviour of IHF-DNA interactions using advanced techniques such as single-molecule fluorescence or NMR spectroscopy, providing a more comprehensive understanding of the conformational dynamics and kinetics of IHF-DNA interactions.

Hypotheses suggest that a well-regulated mechanism is imperative for the successful integration of the phage genome, ensuring its genetic continuity. Given the coexistence of IHF and Holliday junction in a single process and alongside evidence of specific and non-specific protein-DNA binding, the possibility arises of a regulatory role of IHF on Holliday junction. Based on recent findings regarding IHF’s involvement in bacterial biofilms containing HJ (Devaraj *et al*, 2019) and its promotion of homologous recombination in Pseudomonas putida (Mikkel *et al*, 2020), we embarked in investigating the molecular-level binding of IHF to DNA four-way junctions. In this study, we explored IHF’s binding to these junctions in presence of the consensus sequences H’ and H1, utilizing microscale thermophoretic binding assays. Our results indicate that IHF exhibits high affinity for junctions, suggesting that the protein binds to the central region of the junction. Circular dichroism spectroscopic studies imply that the protein induced conformational change to DNA four-way junctions. However, the single molecule FRET experiments unequivocally demonstrate that IHF binds preferentially to the open junctions with high affinity and stabilizes them in that particular conformation.

## Results

### Comparative binding of DNA four-way junctions by WtIHF and ScIHF in absence and presence of MgCl_2_

The attP site comprises two distinct regions, H’ and H2, each possessing a dA+dT rich element. This specific element is effectively protected from nuclease digestion due to its interaction with IHF protein. However, the corresponding segment within the H1 site lack a dA+dT rich element, rendering it susceptible to nuclease cleavage in presence of IHF (Craig & Nash, 1984; Yang & Nash, 1989). This observation suggests that both elements-the dA + dT-rich region and the core consensus determinant-are essential for IHF to bind effectively to the H’ site. On the other hand, the H1 site, without an upstream dA+dT-rich region, can still facilitate IHF binding solely based on the presence of the core consensus determinant (Hales *et al*, 1994).

Binding studies were conducted to assess the affinity of the two proteins WtIHF and ScIHF to the two specific DNA sequences H’ & H1. The binding affinity was first tested at 0mM MgCl_2_ in which the junction adopts an open conformation and it was found that binding constant (K_d_) of WtIHF for H’-HJ and H1-HJ junctions are 2.8±0.5 nM and 11±1.6 nM (Figure 1a) respectively. For ScIHF, K_d_ values for H’-HJ and H1-HJ are 2.86±0.73 nM and 4.22±1.16 nM respectively (Figure 2d). No significant difference is observed for the between the two proteins, indicating that the dA+dT-rich region is not indispensable for binding of either protein to the H’ or H1 sequence. Further K_d_ values at 0mM MgCl_2_ for all combinations of junction and protein were within the range of a few nano molars, indicating high affinity and tight binding. Subsequently, binding affinities were examined at 2mM MgCl_2_, a concentration closer to physiological conditions. The K_d_ values for WtIHF binding to H’ and H1 junctions (Figure 1b) found to be 12±0.89 nM and 7.75±1.1 nM respectively. Similar values were obtained for ScIHF at 3.38±0.34 nM and 5.42±1.17 nM for H’ and H1 sequence harbouring junctions (Figure 1e). Furthermore, binding assays were conducted at higher MgCl_2_ concentration of 50mM. For WtIHF with H’ and H1 junctions, the K_d_ values (Figure 1c)were determined as 13.6±2.21 nM and 7.09±0.93 nM, respectively. The corresponding values for ScIHF were2.26±0.82 nM and 5.96±0.67 nM (Figure 1f).

**Figure 1:**
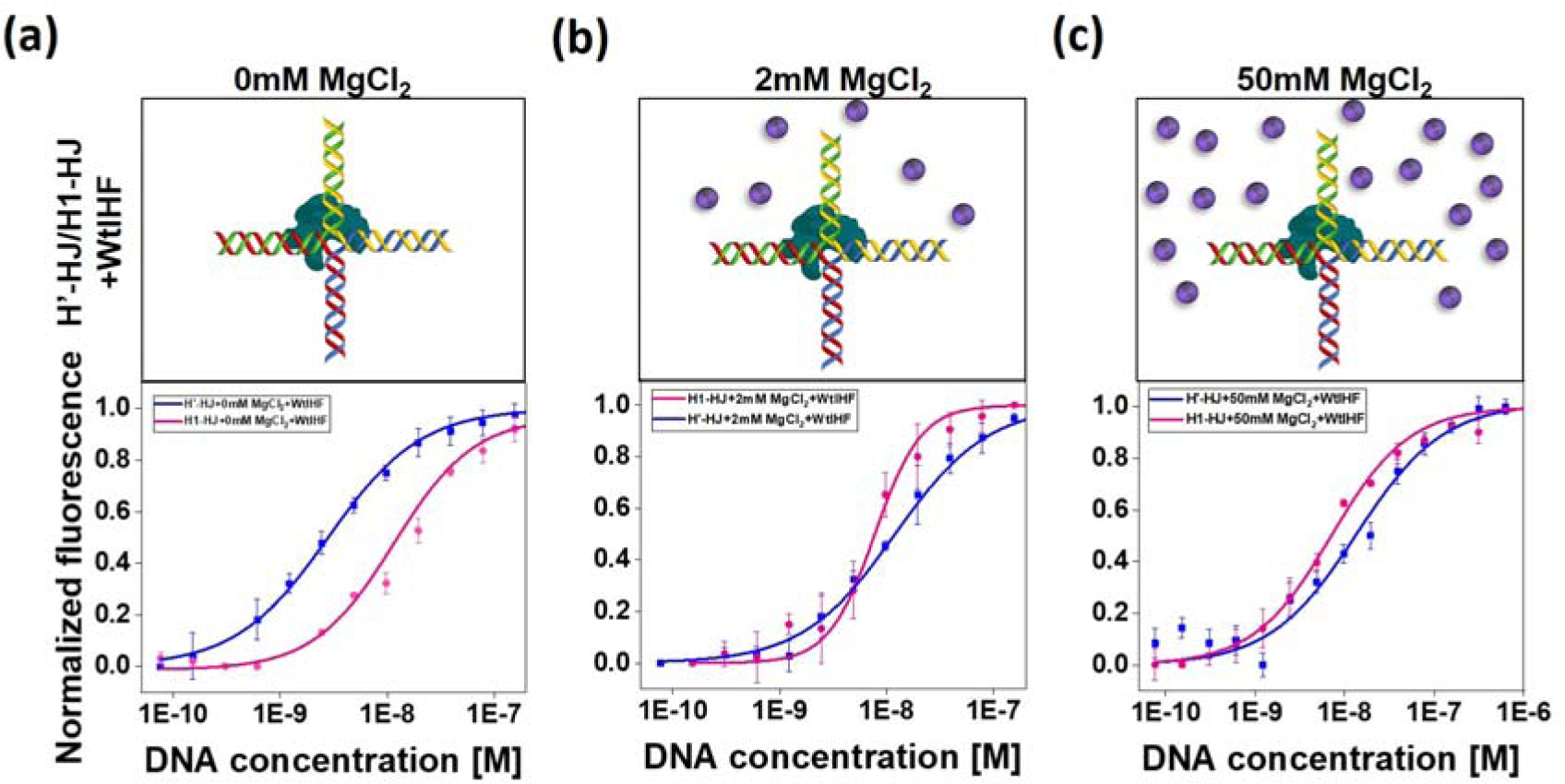
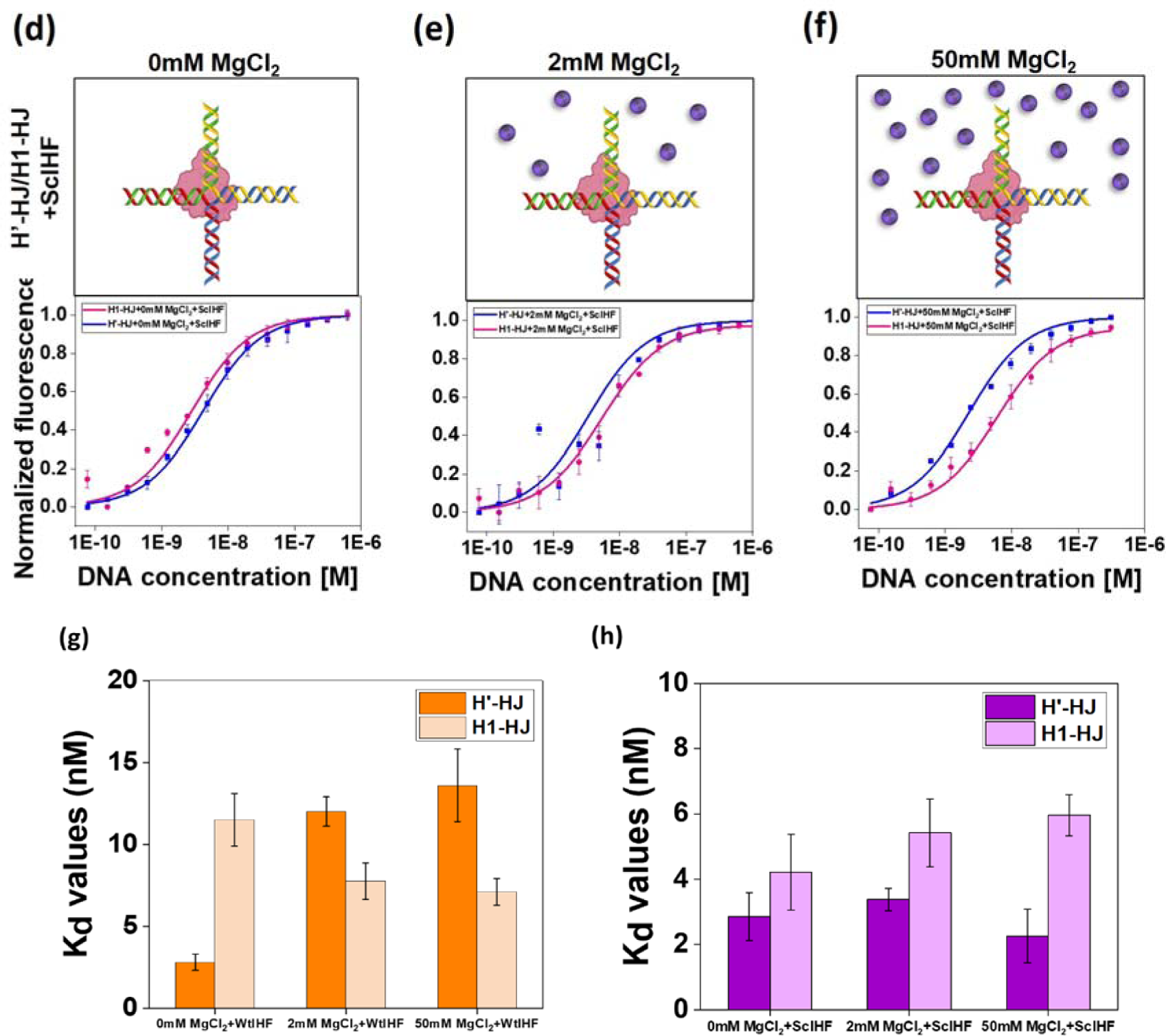
Interaction studies of Holliday junction (H’ & H1) and proteins (WtIHF & ScIHF) by MST. Dissociation constants of the interactions were obtained from the related binding curves. Titration of unlabeled H’-HJ to labelled WtIHF and ScIHF are shown in (•) whereas that of unlabeled H1-HJ is shown in (▪). Binding curves were generated and fitted with Hill equation for the two DNA constructs with the two proteins under different buffer conditions as shown in figure. All data represent the mean (SD) of three independent measurements. (a) Cartoon representation and binding curve of WtIHF to H’, H1 junctions at 0mM MgCl_2_, (b) 2mM MgCl_2_ and (c) 50mM MgCl_2_ (d) Cartoon representation and binding curve of ScIHF to H’, H1 junctions at 0mM MgCl_2_, (e) 2mM MgCl_2_ and (f) 50mM MgCl_2_.

**Figure 2:**
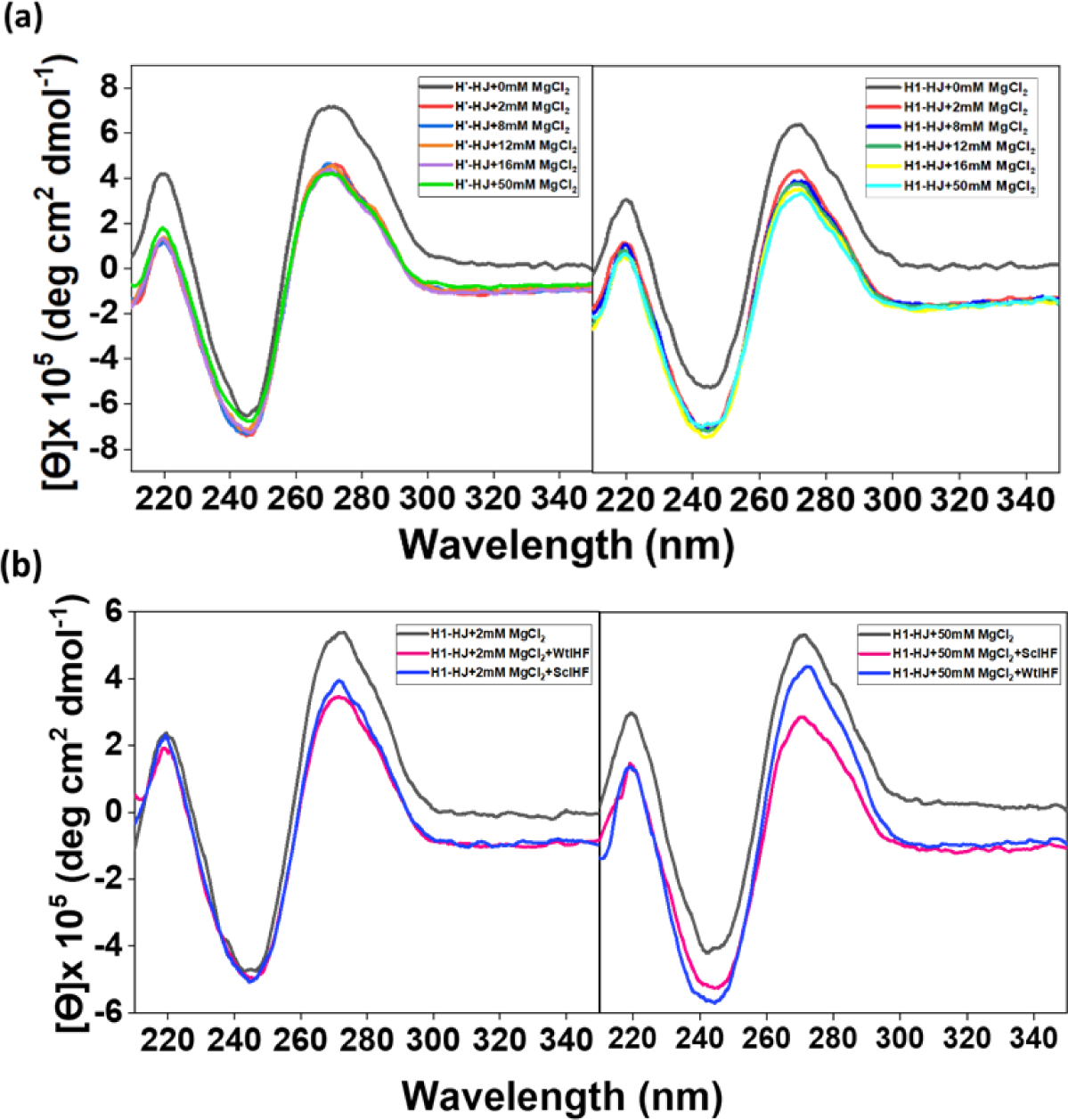
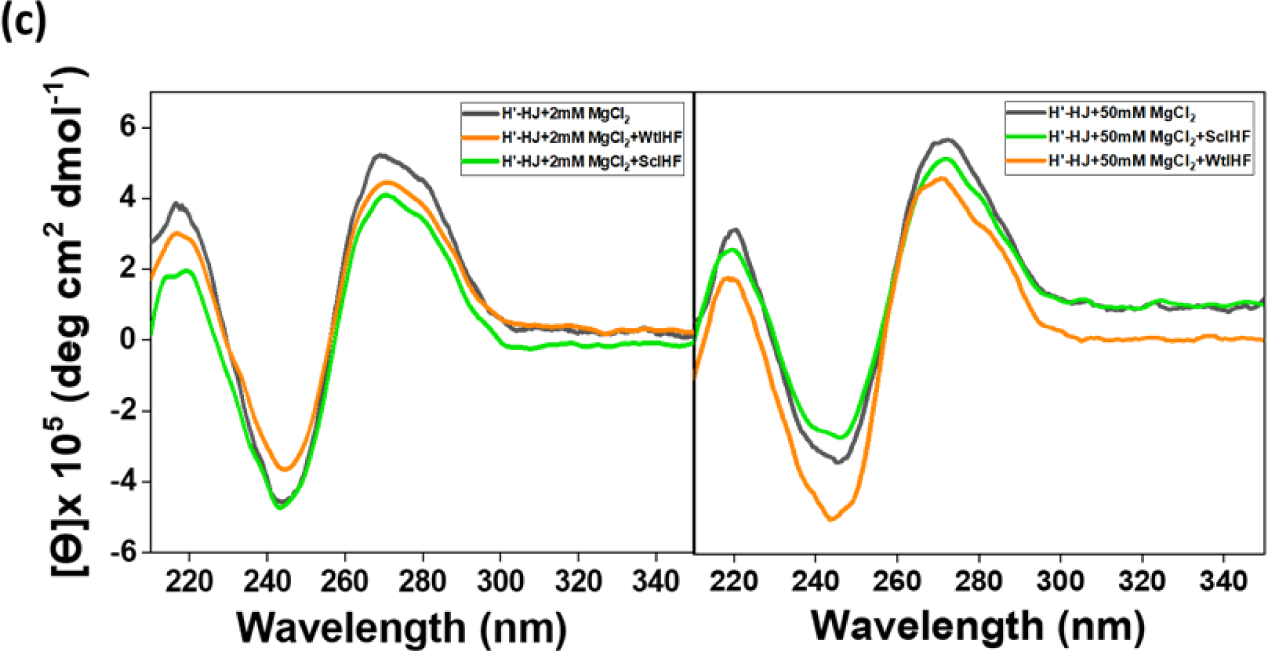
Circular dichroism spectroscopy of Holliday junction dynamics in varied concentrations of Magnesium and in presence of proteins WtIHF & ScIHF. (a) Effect of MgCl2 on the CD spectra of Junctions H’-HJ and H1-HJ. CD spectra overlayed of both junctions with MgCl2 titrated in from 0 mM – 50 mM. The spectra looks similar for both junctions with the dip at 271 nm being most significant from 0 to 2 mM and less significant there onwards. (b) Junction dynamics of H’-HJ and H1-HJ upon addition of 5μM WtIHF and ScIHF.

From the above obtained values, several conclusions can be drawn. First, there is no significant change in binding affinity for either proteins in absence or presence of up to 50mM MgCl_2_. The experiments were conducted in synchronization with our single molecule experimental conditions Across all binding experiments, K_d_ values remained consistently within nano molar ranges, suggesting that the addition of 50mM magnesium did not disrupt or affect protein binding. Furthermore, despite the absence of dA + dT rich regions in our junction construct, no major differences in binding affinities between WtIHF and ScIHF for the H’ and H1 construct were observed. Finally, the low K_d_ values are indicative of tight binding (Figure 1g & h).

### Circular Dichroism analysis of the effect of Holliday junction conformational dynamics

Scientific advancements have unveiled the ability of Holliday junctions (HJ) to adopt two distinct conformations: the Open-X and the Stacked-X forms (Hyeon *et al*, 2012; Joo *et al*, 2004; Grainger *et al*, 1998). An intriguing hypothesis emerged, suggesting that circular dichroism (CD) could serve as effective tool to distinguish between these conformations. in a prior study, the effects of adding MgCl_2_ to a DNA molecule called Cruci3HL, which forms a cruciform structure has been investigated (Hamilton, 2014). Remarkably, the introduction of MgCl_2_led to a significant decrease in the molar ellipticity in the circular dichroism (CD) spectrum. Subsequent additions of MgCl2 exhibited diminishing effect on the CD spectrum. This phenomenon was also observed in Cruci3HL possessing a hairpin loop with in one of the arms (Hamilton, 2014). Similar results were obtained for both mobile and immobile junctions featuring12 bp arms, where the addition of MgCl_2_induced a decline in the molar ellipticity around 275nm. intriguingly the configuration at the junction was influenced by the flanking sequence (Shida *et al*, 1996).

The Stacked-X conformation predominantly manifests itself in the presence of MgCl_2_. To investigate this phenomenon in our junction constructs, numerous CD spectra were acquired during the incremental introduction of MgCl_2_ into solutions containing H’-HJ and H1-HJ. The initial supplementation of 2mM MgCl_2_ resulted in a significant change in the CD spectrum, characterized by a decrease in molar ellipticity at 275 nm (Figure 2a). Successive additions of MgCl_2_ at concentrations of 8, 12, 16, and 50mM induced further, albeit less pronounced, changes in the spectra. Thus, it is reasonable to infer that MgCl_2_ induces structural changes in DNA to a certain extent, likely through interactions with the DNA backbone and nucleobases [28]. Expanding our investigations, we utilized CD as an effective tool to examine if proteins bind to DNA junctions and to track the ensuing structural modifications. Upon introducing both WtIHF and ScIHF to DNA, an observable negative shift in the ellipticity was noted at 271 nm (for both 2 mM and 50mM magnesium concentrations) for both H1-HJ and H’-HJ (Figure 2b & c).

### Characterization of non-migratable Holliday Junction (H’-HJ and H1-HJ) dynamics in presence and absence of Magnesium ions (Mg2+)

Each of the two Holliday junctions, constructed by annealing four single stranded DNA (ssDNA) oligomeric strands, was labelled with Cy3 (donor) and Cy5 (acceptor) fluorophores to monitor conformational dynamics of the junction in absence of a divalent cation (Mg^2+^) and in presence of varying concentrations of Mg^2+^. The sequence and design of the constructs, as shown in (Table 1, Figure 3), were carefully chosen to ensure that both the junction are non-migratable and harboured a distinct protein binding sequence (H’ or H1) on one of its component strands. The central sequence, comprising the base pairs at the junction or crossover point was identical for both constructs. Cy3 (donor) and Cy5 (acceptor)were attached to two adjacent helical arms to probe the dependence of the junctions on divalent metal ions to undergo conformer transition between the stacking conformers.

**Figure 3:**
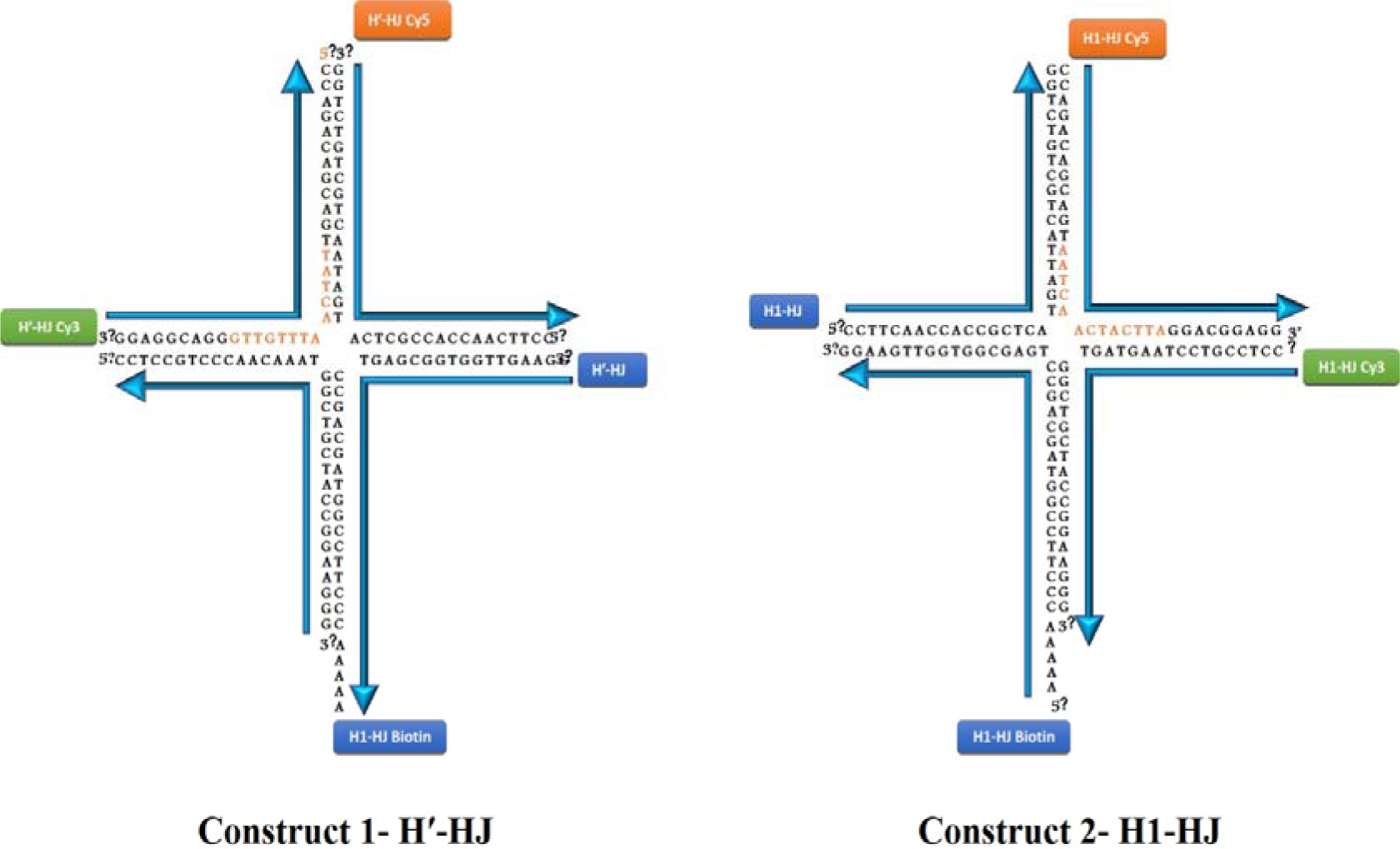
Pictorial representation of the two DNA four-way junctions used in the study. Protein binding sequences (H’ & H1) are shown in orange. The four strands are labelled as shown in the picture with labelling of Cy3 and Cy5 in adjacent arms so as to capture conformational change under different experimental conditions. Each arm of the junction consists of 17 base pairs.

**Table 1:**
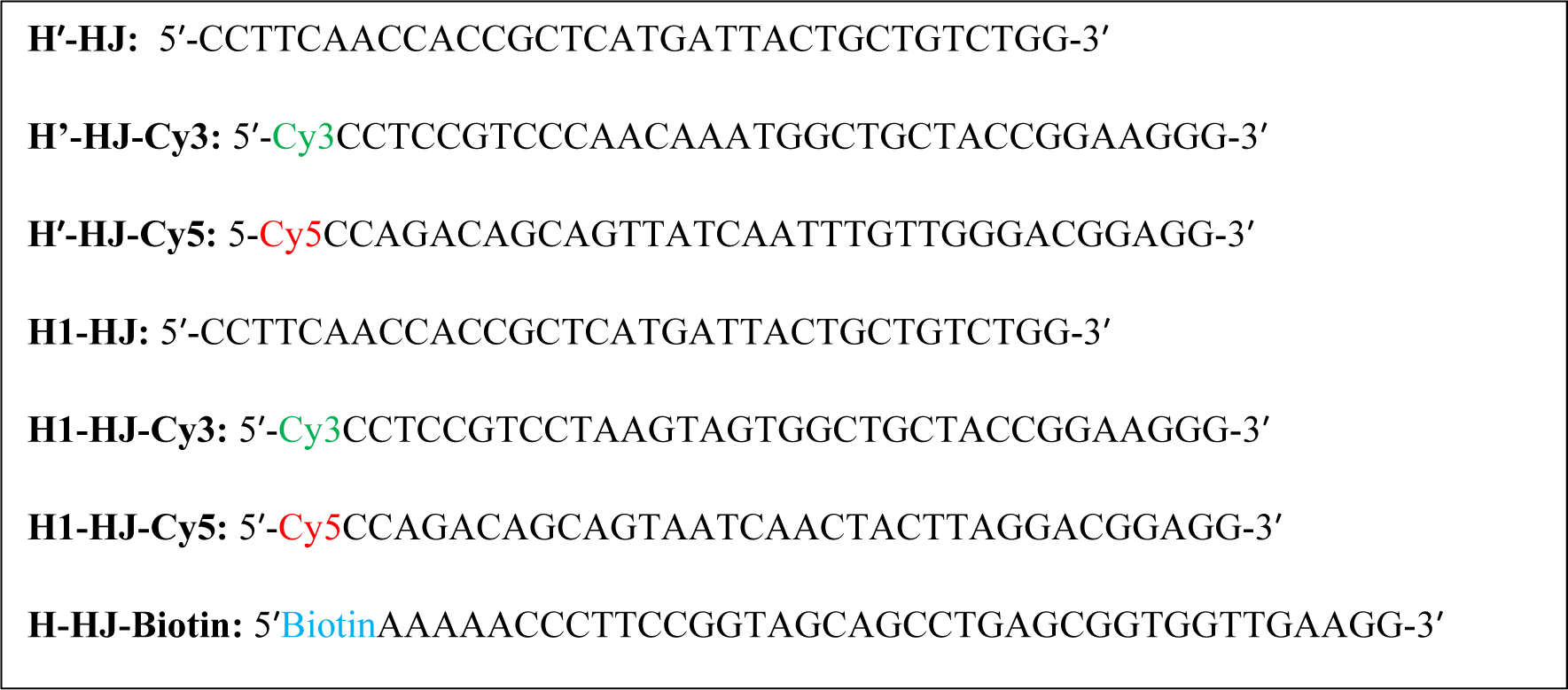
Oligonucleotide sequences used to construct the two junctions, H’ and H1.

The occurrence of FRET between donor and acceptor fluorophores, based on the physical distance between them, indicated different conformational states adopted by the junctions under different salt concentrations. In absence of any divalent cation, junctions are known to exist in the open conformation with a four-fold symmetry wherein each of its four helical arms extends at nearly 90° angle to its adjacent arm (Promisel *et al*, 1989; Duckett *et al*, 1988). The top tier of Figure 4c & 4d displays fluorescence signals from donor and acceptor as a function of time, along with the corresponding FRET efficiency trace of an individual H’-HJ molecule in presence of T50 Buffer (10mM Tris-Cl, pH 8.0, 50mM NaCl). The FRET efficiency trace reveals no change in FRET efficiency till photobleaching. The corresponding ensemble histogram (Figure 4a) has a single Gaussian peak with an apparent FRET efficiency (E_FRET_) of 0.38. We assigned this FRET state to the open-X conformation of the junction. Furthermore, all of the ∼200 molecules analysed were found to reside in the open conformation, as portrayed by the individual histogram plot generated by superimposing individual histogram of all 200 molecules (Figure 4b).Similar results were obtained with the junction H1-HJ (Figure 4e-h).

**Figure 4:**
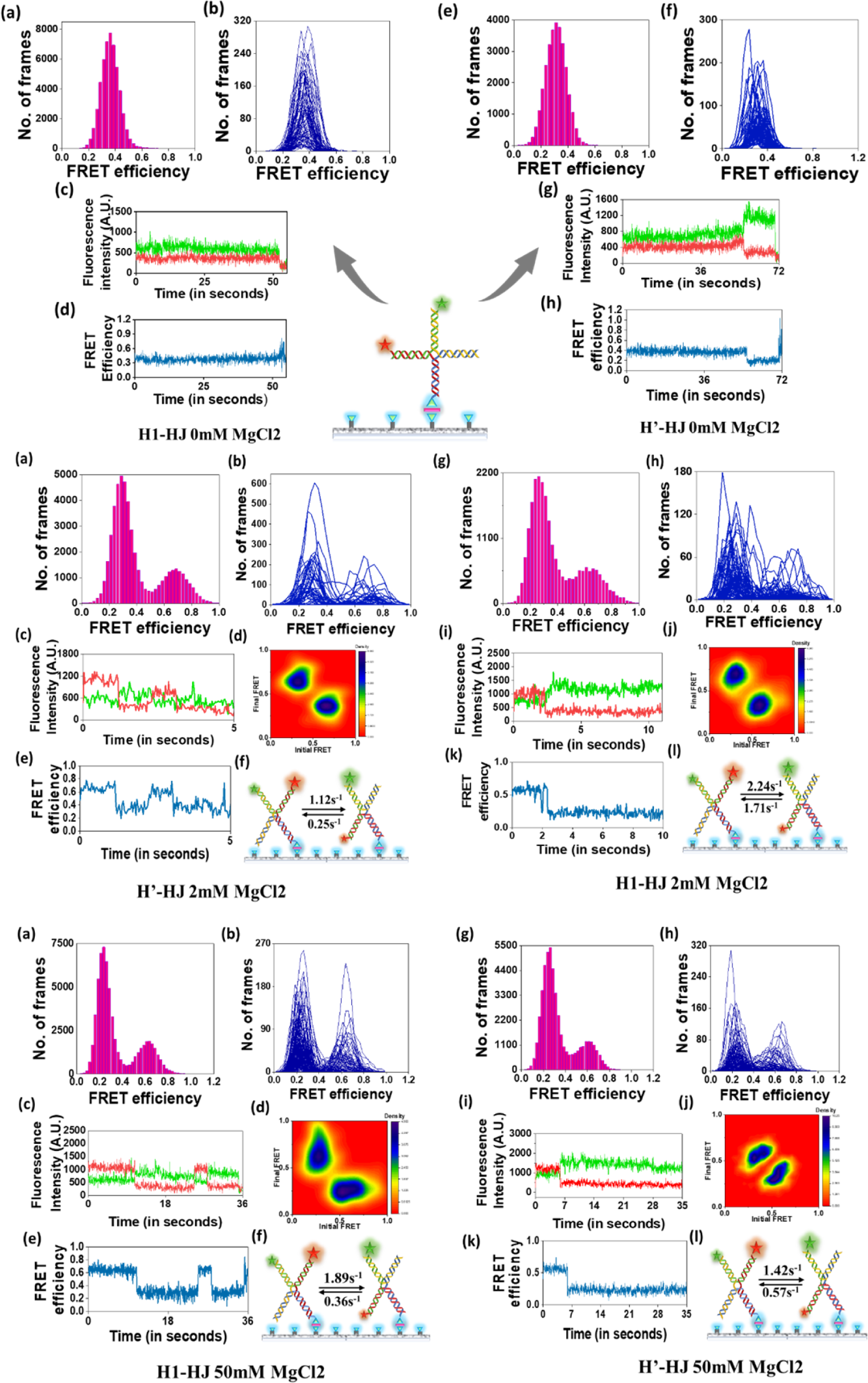
Top at 0mM MgCl_2_: (a,b) Ensemble histogram and Individual histogram showing a single FRET state at 0.38, (c,d) Representative time trace of fluorescence intensity and FRET efficiency of an individual molecule for the junction H’-HJ; (e-h) Ensemble histogram, individual histogram, Representative time trace and FRET efficiency plot for H1-HJ. Middle at 2mM MgCl_2_: (a-f) Ensemble and individual histogram, representative time trace and FRET efficiency plot, Transition density plot TDP, schematic diagram of transition rate between isoⅠ and isoⅡ for junction H’-HJ. (g-l) Ensemble and individual histogram, representative time trace and FRET efficiency plot, Transition density plot TDP, schematic diagram of transition rate between isoⅠ and isoⅡ for junction H1-HJ. Bottom at 50 mM MgCl_2_: (a-f) Ensemble and individual histogram, representative time trace and FRET efficiency plot, Transition density plot TDP, schematic diagram of transition rate between isoⅠ and isoⅡ for junction H’-HJ. (g-l) Ensemble and individual histogram, representative time trace and FRET efficiency plot, Transition density plot TDP, schematic diagram of transition rate between isoⅠ and isoⅡ for junction H1-HJ.

In the context of stacked X-isomers, we observed two distinct conformations, X-isomer-I and X-isomer-II. In X-isomer-I, DNA strands are arranged in a specific stacked configuration, forming an X-shaped junction., while isomer-II exhibits a different stacked arrangement, leading to an alternative X-shaped conformation. Previous research has established that the distribution of isomers in Holliday junctions (HJs) depends on the preferential stacking of nucleotide bases at the junction (Grainger *et al*, 1998). The specific arrangement of nucleotide bases and their interactions play a crucial role in determining the stability and conformational preferences of the HJ (Hays *et al*, 2003; Khuu *et al*, 2006). In stacked X-isomers (isomer-I and isomer-II), nucleotide bases at the junction engage instable stacking interactions, resulting in the characteristic X-shaped conformation. The sequence and nature of the nucleotides can influence the stability of these stacked conformations, with certain nucleotide sequences favouring stronger stacking interactions, thus biasing the equilibrium between the two isomers. Conversely, in the open unstacked conformation, the nucleotide bases at the junction do not engage in stable stacking interactions, leading to a more extended and flexible structure. The absence of strong stacking interactions allows the HJ to adopt an open conformation. Various factors influence the preferential stacking of nucleotide bases is influenced by, including the sequence composition of the DNA arms, the presence of specific ions (such as Mg^2+^), and the binding of proteins or ligands. These factors collectively modulate the stability of the different isomers, leading to conformational changes and dynamic behaviour of the HJ.

We attempted to study the effect of a divalent cation (Mg^2+^) at a physiological concentration of 2mM (Buffer1; 10mM Tris-Cl, pH 8.0, 50mM NaCl, 2mM MgCl_2_) on junction dynamics. Both the ensemble as well as individual histogram plots (Figure 5a-b, middle) have two peaks corresponding to FRET efficiency values of 0.22 and 0.66. We assigned the low FRET with E_FRET_ = 0.22 as isoⅠ and the high FRET state with E_FRET_= 0.66 as isoⅡ for Hʹ-HJ. These two states represent the inter conversion of Hʹ-HJ between the two stacked conformers with a clear bias towards the low FRET state (isoⅠ).The time trace and FRET efficiency trace of a single representative molecule is also shown (Figure 4c-d).The transition density plot (TDP) also shows conformational dynamicity between two FRET states (Figure 4e).Time traces of all such molecules exhibiting transition between the two FRET states (high and low) were assigned to two-state HMM (Hidden Markov Model) analysis and the output generated in the form of a transition matrix was used to derive values for temporal kinetic rates of transition. The rate of transition from isoⅠ to isoⅡ (k_I→II_)was found to be 0.25s^-1^ whereas k_II→I_ was obtained as 1.12s^1^(Figure 5f). The mean dwell time (time spent by a molecule in a particular conformational state) calculated manually for H’-HJ in isoⅠ (low FRET state) is τ_1_=4s and in isoⅡ (high FRET state) is τ_2_=0.89s. Difference in free energy (ΔG) between the two stacking conformers (isoⅠ and isoⅡ) was calculated by -RTln(k_I→II_/k_II→I_)and its value was obtained as 3.6kJ mol^-1^ which is consistent with the value (∼3.0kJ mol^-1^) reported in literature (Mckinney *et al*, 2002) The junction H1-HJ also exhibits similar trend with transition from isoⅠ to isoⅡ (k_I→II_) obtained as 1.714s^-1^ whereas (k_II→I_)was obtained as 2.248s^1^. The mean dwell time for states isoⅠ and isoⅡ are found to be 1.91s and 0.44s respectively(Figure 4g-l).

**Figure 5:**
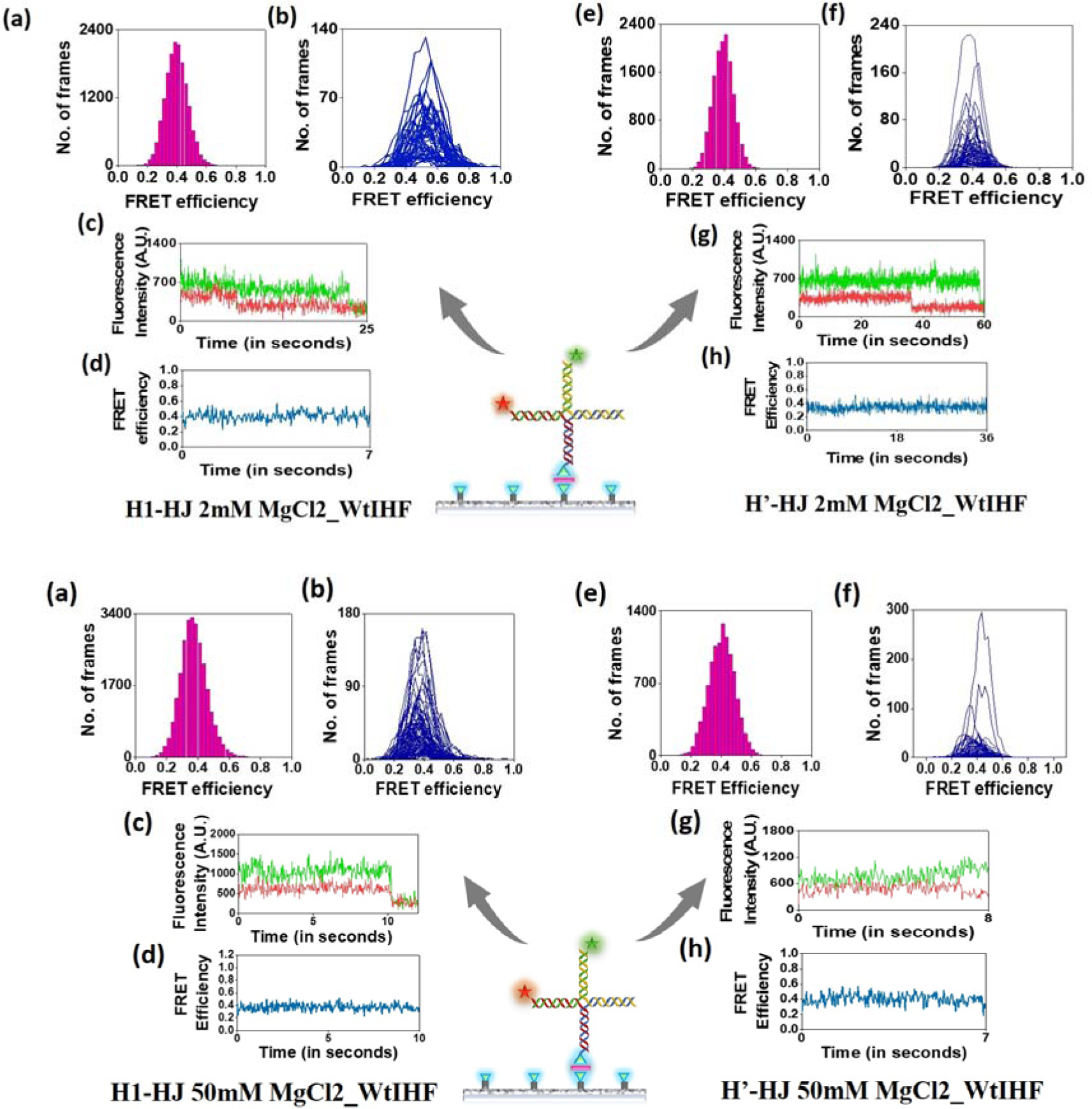
Top at 2mM MgCl_2_ with WtIHF: (a-f) Ensemble and individual histogram, representative time trace and FRET efficiency plot, for junction H’-HJ. (e-h) Ensemble and individual histogram, representative time trace and FRET efficiency plot for junction H1-HJ. Bottom at 50 mM MgCl_2_ with WtIHF: (a-d) Ensemble and individual histogram, representative time trace and FRET efficiency plot, for junction H’-HJ. (e-h) Ensemble and individual histogram, representative time trace and FRET efficiency plot for junction H1-HJ. In almost all cases a single FRET state with FRET value of 0.38 is obtained.

The experimental condition was further extended to a higher concentration of Mg^2+^ to check if increasing the cationic concentration had any effect on the population distribution or longevity of a specific state in which molecules existed. For this, experiments were performed with Buffer 2 (10mM Tris-Cl, pH 8.0, 50mM NaCl, 50mM MgCl_2_). Ensemble and individual histograms of the junction H’-HJ again depict two distinct FRET states (Figure 5a-b, bottom), a low FRET state at 0.23 and a high FRET state at 0.63. Fluorescence intensity and FRET efficiency plots as well as TDP shows conformational exchange between the two states (Figure 5c-e). The low FRET state corresponds to isoⅠ while the high FRET state is indicative of isoⅡ. Rate of conversion from isoⅠ to isoⅡ (k_Ⅰ→Ⅱ_) equals 0.574s^-1^ and from isoⅡ to isoⅠ (k_Ⅱ→Ⅰ_) is 1.425s^-1^. Mean dwell times calculated for molecules existing in isoⅠ (τ_1_) was found to be 1.74s whereas for isoⅡ (τ_2_) was 0.7s. Free energy difference was calculated as 2.3kJ mol^-1^ which is close to reported value (∼3kJ mol^-1^). Experiments with the second construct (H1-HJ) were also performed with 50mM Mg^2+^ which yielded similar results (Figure 4g-j). Rate of conversion from isoⅠ to isoⅡ (k_Ⅰ→Ⅱ_) equals 0.36s^-1^ and from isoⅡ to isoⅠ (k_Ⅱ→Ⅰ_) is 1.897s^-1^. Mean dwell times calculated for molecules existing in isoⅠ (τ_1_) is found to be 2.77s whereas for isoⅡ (τ_2_) is 0.52s.No significant difference was found in conversion rates at 2mM and 50mM MgCl_2_. In conclusion, both junction constructs exhibited typical behaviour under the influence of the divalent cation magnesium. In the absence of magnesium, with only a small amount of sodium, the junctions remained in an open state. However, upon the addition of magnesium to the medium, the junctions alternated between two stacking conformers.

### Wild-type Integration Host Factor alters the conformational dynamics of the Holliday Junction

Wild-type Integration Host Factor (WtIHF) is categorized as an architectural protein, a crucial class of DNA-binding proteins that play a pivotal role in bringing distant genomic sites into proximity, often involving DNA bending. This architectural function further contributes to the organization of chromatin into higher-order structures (Yoshua *et al*, 2021). In the context of DNA duplex bending, WtIHF is well-known for facilitating essential cellular processes, including site-specific recombination, replication, transcription, and DNA compaction. Recent SmFRET studies (Purkait *et al*, 2021), atomic force microscopy investigations, and molecular dynamics simulations have shed light on the multimodal bending of DNA duplexes induced by IHF (Yoshua *et al*, 2021). IHF-induced DNA bending has been a well-established concept stemming from structural studies. However, IHF’s abundance as a Nucleoid Associated Protein (NAP) in prokaryotic cells, with approximately 12,000 molecules per cell in the exponential growth phase and 55,000 molecules per cell in the early stationary phase (Azam *et al*, 1999), prompts questions about its roles beyond structural alterations to DNA duplexes. It is firmly established that IHF is a pivotal protein supporting site-specific recombination of phage DNA in E. coli, a process involving the formation of a Holliday junction for genetic material exchange.

Based on prior reports on DNA bending by IHF, it was hypothesized that IHF, upon binding, would bend the DNA arms of the junction, potentially altering the distance between fluorophores, thus affecting FRET efficiencies. These changes would be reflected in single-molecule traces and histograms. To observe any modifications in junction conformation resulting from protein binding, it was necessary to stabilize the junction in a specific conformation. To achieve this, the protein’s effect was examined on H’-Holliday Junction (HJ) in T50 Buffer (No Mg^2+^). Contrary to expectations, no change in FRET state was observed upon protein binding. A single intermediate FRET state at 0.38 was obtained (supplementary figure 2a-b). Representative time traces and corresponding FRET efficiency traces (supplementary figure 6c-d) of single molecules showed no alterations in FRET upon protein binding. Similar results were obtained with the second construct, H1-HJ (supplementary figure 2e-h). These findings led to further investigation into the potential effects of the protein on the junction’s dynamics.

Previously, single-molecule Förster resonance energy transfer (smFRET) and ensemble fluorescence techniques were used to study the binding interaction between RuvA and Holliday junction. The results revealed that at physiologically relevant concentrations of Mg2+ ions, the binding of RuvA was found to effectively halt the conformational dynamics of the Holliday junction (Gibbs & Dhakal, 2018). So, SmFRET experiments of the two constructs (H’-HJ and H1-HJ) were performed under a physiologically relevant Mg2+ concentration of 2mM in presence of WtIHF. Whereas in absence of protein, for H’-HJ we observed two peaks at 0.22 and 0.66, a single FRET state with a peak at 0.4 (Figure 5a-b, middle) appears in presence of WtIHF. Figure 6c-d shows representative time trace and FRET efficiency trace of a single molecule in presence of WtIHF. The long representative time traces are indicative of stable protein binding. For H1-HJ, WtIHF yielded identical results (Figure 5e-h). Overall, the emergence of a single FRET peak around 0.38 unequivocally demonstrates that WtIHF clamps the four-fold symmetrical open conformation of the Holliday junction, thereby halting the conformational interconversion between the two stacked-X conformers. Additionally, SmFRET time traces show a continuous static FRET throughout the observation time till photobleaching that indicates formation of a highly stable WtIHF-HJ complex. While at 50mM Mg^2+^ the junction presented itself in two FRET states of 0.22 and 0.64 corresponding to the two isomeric forms. Interestingly, when we performed SmFRET analysis of the Holliday junction in presence of WtIHF under the same buffer conditions (Buffer 2), a single peak at 0.4 appeared along with complete disappearance of the two peaks at 0.22 and 0.64 respectively. The ensemble and individual histogram plots (Figure 5a-b, bottom) clearly depict the changes in peak position. Representative time trace of fluorescence signals from donor and acceptor labelled H’-HJ upon protein binding and FRET efficiency trace (Figure 5c-d) are shown which indicate that the junction existed in a single FRET state around 0.4 till photobleaching. Similar results were established upon protein binding to our second construct (H1-HJ) which harboured a different protein binding sequence, H1 (Figure 5e-h).This indicates the existence of a single peak at 0.38 with loss of the two peaks at 0.2 and 0.62 upon protein binding. Fig 3 depicts time traces and FRET efficiency trace of H’HJ at 50mM MgCl_2_ in presence of WtIHF. Reportedly, this is the first SmFRET-based visualisation of the manipulation of Holliday junction conformational dynamics by an architectural protein at the single molecule level.

**Figure 6:**
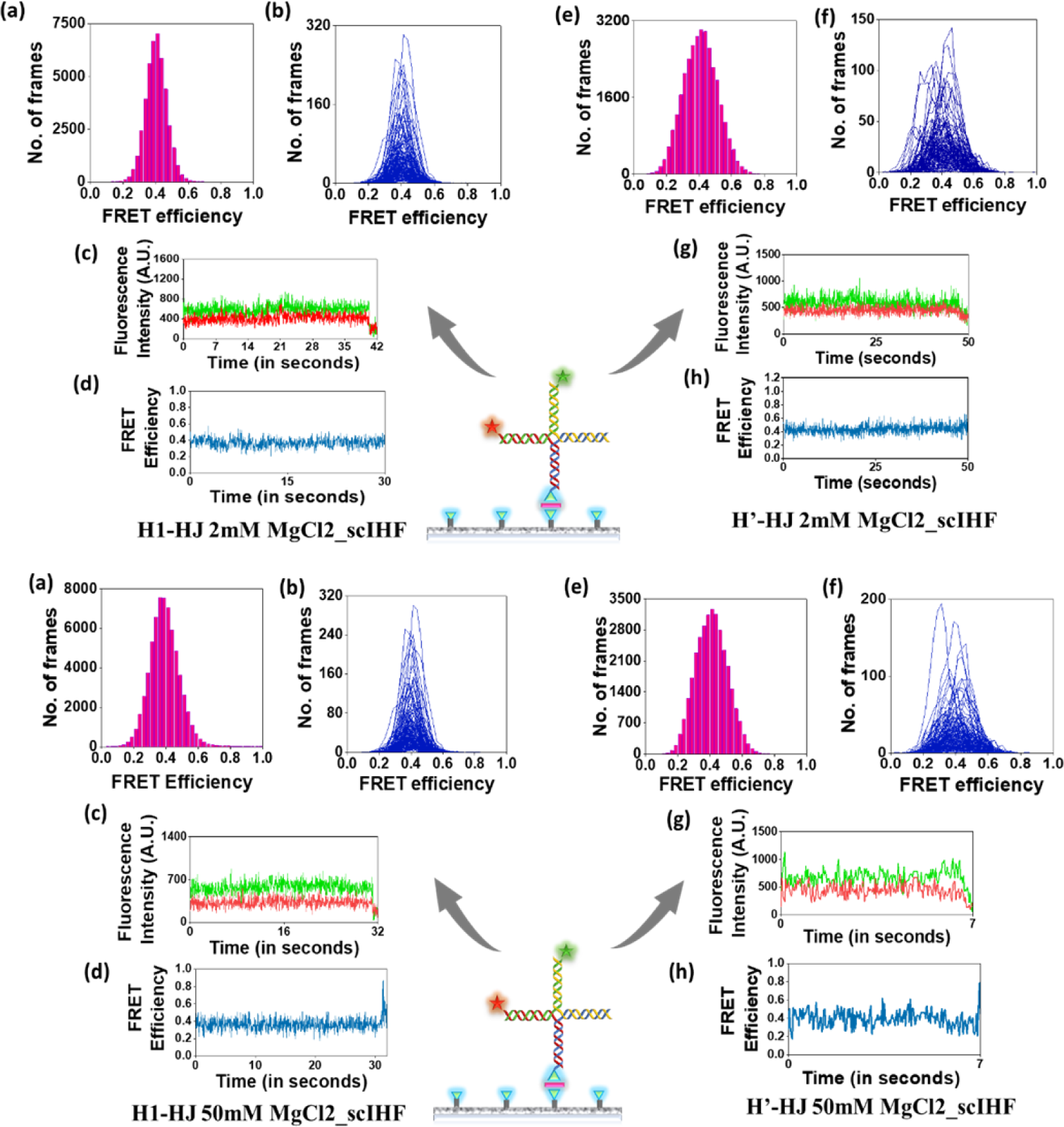
Top at 2mM MgCl_2_with ScIHF: (a-f) Ensemble and individual histogram, representative time trace and FRET efficiency plot, for junction H’-HJ. (e-h) Ensemble and individual histogram, representative time trace and FRET efficiency plot for junction H1-HJ. Bottom at 50 mM MgCl_2_with ScIHF: (a-d) Ensemble and individual histogram, representative time trace and FRET efficiency plot, for junction H’-HJ. (e-h) Ensemble and individual histogram, representative time trace and FRET efficiency plot for junction H1-HJ. In almost all cases a single FRET state with FRET value of 0.4 is obtained.

### Comparative junction dynamics by an engineered variant of IHF, the Single Chain Integration Host Factor (ScIHF)

Numerous biochemical studies reported that single chain IHF, a more stable variant, exhibits comparable activity to WtIHF on duplex DNA (Bao *et al*, 2004). This prompted us to check if the protein’s activity on junction is aligns with that of WtIHF. ScIHF activity was first studied on H’-HJ (supplementary figure 3a-d) and H1-HJ in absence of Mg^2+^ (only T50 Buffer). In presence of ScIHF, a single gaussian peak is observed at 0.4 for H’-HJ and at 0.38 for H1-HJ (supplementary figure 3e-h).

At 2mM Mg^2+^ for H’-HJ in presence of ScIHF, the results were consistent with those reported for WtIHF activity. The two FRET peaks at 0.2 and 0.6 disappeared, replaced by a single peak at 0.4 (Figure 6a-d, middle). Similar results were observed for H1-HJ at 2mM Mg^2+^ with ScIHF, where an individual peak corresponding to a FRET state of 0.39 is evident (Figure 6e-h). Time trace and FRET efficiency trace indicate formation and stabilization of open form of the junction upon bounding to ScIHF. The protein activity was next checked at 50 mM MgCl_2_ for both constructs. For H’-HJ, the two peaks (at 0.23 and 0.63) observed in the absence of any protein disappeared, giving rise to a new single peak at 0.41 (Figure 6a-d, bottom),suggesting the existence of the open conformation of the junction induced upon protein binding. In case of H1-HJ, a similar trend was observed, with loss of the two peaks at 0.22 and 0.64, replaced by an individual peak at 0.37 (Figure 6e-h). Collectively, for both the constructs, ScIHF upon binding to the junction caused it to open up and stabilized the open conformation. in summary, it is unequivocal that, it is ScIHF does not exhibit any significant difference in terms of altering junction dynamics compared to WtIHF and both WtIHF and ScIHF induce interconversion between the stacked-X isoforms upon binding.

## Discussion

The emergence and subsequent diversification of life over the past 3.8 billion years, progressing from unicellular organisms to multicellular entities, owe their feasibility to the presence and dissemination of genetic information contained within nucleic acids, namely DNA or RNA. Nevertheless, genomic DNA, prevalent across most life forms, constitutes a substantial biomolecule susceptible to various forms of stress that may affect its structural and functional attributes. In response to these challenges, nature has devised a multitude of mechanisms for repairing alterations in DNA structure or sequence, thereby restoring the normal functionality of this biomolecule. A significant portion of DNA repair pathways involves the Holliday junction, whose formation, maintenance, and resolution are all pivotal for sustaining genetic stability and fostering genetic diversity.

The initial phase of our study is focused on monitoring junction dynamics under varying concentrations and species of cations. It is found that the concentration of a divalent cation (Mg2+) does not significantly influence the conformer bias of a junction. Our experimental findings consistently show that, for both Mg2+ concentrations tested (2mM and 50mM), the two junction constructs predominantly favoured the isoⅠ conformer. This aligns with prior reports (Joo *et al*, 2004). It is noteworthy that previous single-molecule investigations into the Holliday junction have explored the effect of cation concentration in conjunction with the type of ionic environment on conformer transitions. However, our choice of cationic concentrations in all experimental buffers closely mimics or approximates physiological levels. Consequently, we did not investigate junction dynamics at extremely high ionic concentrations, as our primary objective was to monitor protein activity and its associated changes in the junction under conditions resembling physiological levels.

This rationale also underlies our inclusion of 50mM Na+ in all buffers relevant to single-molecule experiments, given that intracellular sodium concentrations are approximately 15mM. While our chosen sodium ion concentration slightly exceeds the actual intracellular value, it is well-established that significant alterations in junction dynamics due to the presence of monovalent ions occur only at much higher concentrations (∼1M Na+). Our experiments with T50 buffer, devoid of magnesium, reinforce the fact that in the absence of divalent cations, electrostatic repulsion from negatively charged phosphates supersedes the free energy obtained from helical stacking of the junction arms.

Subsequently, we delved into the novel potential of SmFRET to elucidate the manipulation of conformational dynamics within the junction through architectural proteins, namely WtIHF and ScIHF. This dynamically adaptable IHF-bound junction likely facilitates the binding of other junction-binding proteins, such as resolvases. In contrast, the IHF structural homolog HU binds to four-way junctions in a 2:1 ratio and stabilizes the stacked conformation. Given the abundance of these proteins in the cell and their contrasting interactions with the junction, the competition between IHF and HU binding suggests that a delicate balance between junction opening and stacking may be mediated by these proteins. IHF, by inducing the formation of a dynamic open conformation, possibly facilitates branch migration, whereas the stacked isoform induced by HU may be necessary for junction resolution. Consequently, the interplay between HU and IHF binding may play a crucial role in regulating junction migration and resolution.

Through single-molecule analysis, we have demonstrated how a conventional architectural protein, known to induce varying degrees of sharp bends in DNA upon binding, interacts with and stabilizes the Holliday junction in the open conformation. Our findings further reveal that at physiological concentrations of free Mg2+—2mM or even at a higher concentration of 50mM—both WtIHF and ScIHF can effectively bind and form stable complexes with the junction, indicating unhindered protein-DNA interactions under these magnesium ion concentrations in the environment (Figure 8). In conclusion, our study highlights the capability of an engineered variant of a wild-type protein, ScIHF, to effectively halt junction dynamics.

**Figure 7:**
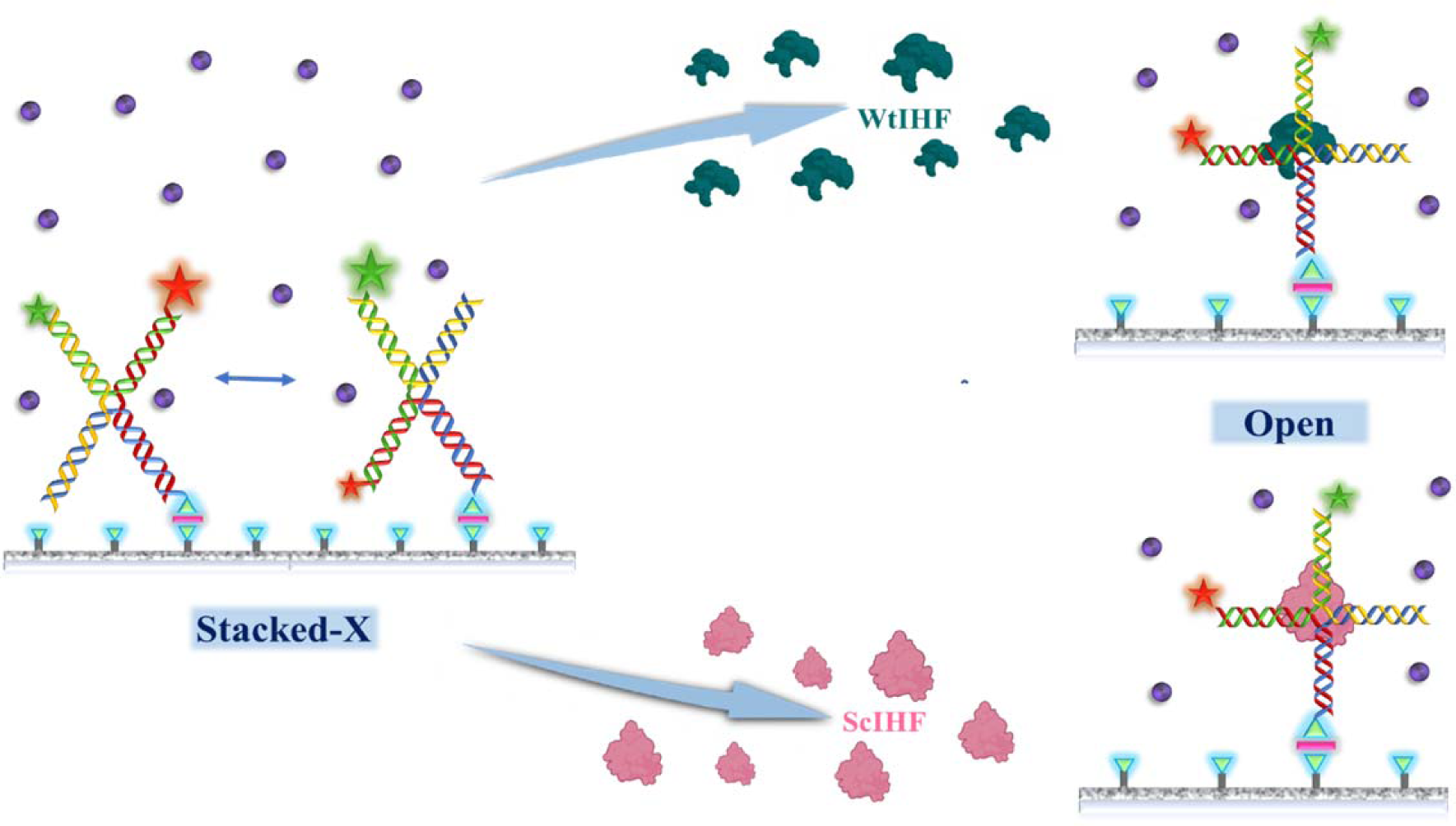
Model representing Holliday junction conformational dynamics in presence of magnesium ions, in which the junction alternates between the two stacked-X forms. The proteins, WtIHF and ScIHF upon binding to the junction halts this conformational dynamicity while inducing and stabilizing the open form of the junction that is favourable for branch migration.

**Figure 8:**
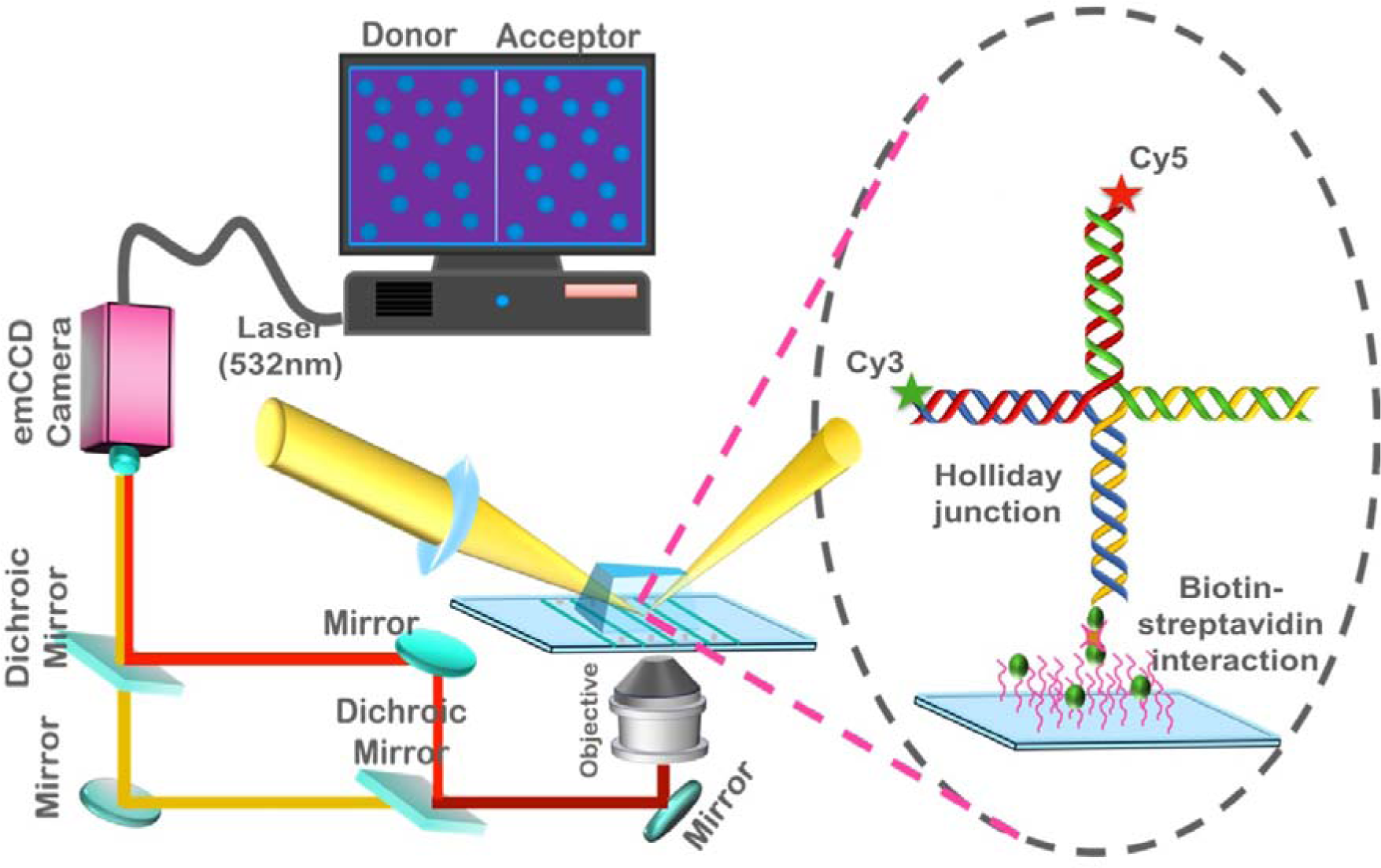
Pictorial representation of Prism-type total internal reflection (PTIR) SmFRET-Single-molecule Forster resonance energy transfer employed in surface tethered experiments in which surface immobilization of Holliday junction is achieved through the strongest non-covalent biotin-streptavidin interaction.

## Materials & methods

### Reagents

The chemical and reagents (analytical grade) used for the smFRET experiments were of analytical grade, purchased either from Sigma-Aldrich (St. Louis, MO) and Merck (Kenilworth, NJ). The standard salts used in formulation of various buffers were procured from either Sigma-Aldrich or Sisco Research Laboratories Pvt. Ltd. (SRL, Mumbai, India). The growth media used for bacterial cell culture were from HiMedia Laboratories. For both protein purification process and single molecule experiments, all the buffers and media were prepared with ultra-purified water from Arium Comfort Series, Sartorius.

### Induced Over-expression of Wild-type Integration Host Factor (WtIHF) and Single-chain Integration Host Factor (ScIHF) in *E.coli*

For the purpose of over-expression, *Escherichia coli* BL21(DE3) cells were transformed with both WtIHF and ScIHF plasmids. Following this, the Cells were grown overnight at 37°C on Luria-Bertani agar (LB-Agar) plates supplemented with Ampicillin. The transformed cells were subsequently cultivated in 1 litre LB broth supplemented with 100μg/ml Ampicillin at 37°C for 3 hours till O.D. reached to 0.6. Following this, The cells were induced with 2mM isopropyl-1-thio-β-D-galactopyranoside (IPTG) and allowed to grow overnight for 14 hours at 20°C. Then the culture media were centrifuged (Heraeus, Biofuge, Stratos, Thermo Fisher Scientific) at 5000rpm for 10 minutes at 4°C and the resulting cell pellet was collected. This pellet was subsequently washed twice with ultra-purified water containing 10% glycerol and then stored at −80°C.

### Purification Of WtIHF and ScIHF

The proteins of interest, both WtIHF and ScIHF have an approximate molecular weight of 22 kilo-Dalton (kDa) and were purified with highest possible purity by employing Immobilised Metal Affinity Chromatography (IMAC). The cell pellet was resuspended in Lysis Buffer (20mM Tris-Cl, pH 8.0, 20mM NaCl, 1mM EDTA, 10% glycerol) to which Protease Inhibitor Cocktail (PIC) purchased from Sigma-Aldrich was added at a dilution of 1:1000. After this, the cells were sonicated (Ultrasonic Processor, Cole Parmer) with 10 cycles of 30 seconds ‘ON’ followed by 30 seconds ‘OFF’ pulses at 20% amplitude.

The lysed cells were centrifuged at 10000 rpm for 40 minutes at 4°C. The supernatant was collected was subsequently collected and loaded on a Ni-NTA column pre-equilibrated with 5 column volume (CV) of chilled Lysis Buffer. The supernatant was passed through the column three times to ensure maximum binding of the desired protein to the beads. The column was then washed with 20 CV of Wash Buffer (20mM Tris-Cl, pH 8.0, 20mM NaCl, 1mM EDTA, 40mM Imidazole) to get rid of any unwanted non-specific proteins bound to the Ni-NTA agarose beads. Thereafter, the proteins (WtIHF or ScIHF) was eluted with 10 ml of Elution Buffer (20 mM Tris-Cl, pH 8.0, 1mM EDTA, 400 mM Imidazole) in fractions of 1 ml each. The flow rate of the column was kept constant at 0.5 ml/min for the loading, washing and elution steps. Presence of protein in eluted samples was checked by Bradford Assay while the purity of the elute fractions was confirmed by Polyacrylamide Gel Electrophoresis (SDS-PAGE, 5% Stacking, 15% Resolving). Thereafter, elute fractions with a single band at 22kDa corresponding to the protein molecular weight was pooled down.

Pooled fractions were then concentrated using 50ml 10kDa Amicon Ultracentrifugal Filter (Merck) by centrifuging at 2500rpm for 5 minutes at 4°C. Concentrated protein was then subjected to single step buffer exchange with Dialysis Buffer (20mM Tris-Cl, pH 8.0, 50mM NaCl) with the help of aPD10 desalting column (GE Healthcare, Sephadex G-25 M matrix) under gravity flow. The resulting elute was then re-concentrated using 10kDa Amicon Ultracentrifugal Filter (Merck) by centrifuging at 2500rpm for 5 minutes at 4°C.

The concentrated protein (WtIHF or ScIHF) were collected and a final qualitative and quantitative check was done by running a 15% SDS-PAGE (supplementary figure 4) followed by determining protein concentration (Absorbance at 280 nm) with the help of Nanodrop Spectrophotometer (Thermoscientific Nanodrop One^c^). Concentrated protein sample was then divided in small aliquots of 50 μl each and utilised for smFRET experiments within 3-4 days of purification to avoid protein degradation or denaturation. The remaining aliquots were stored in −80°C. The whole purification process was performed at 4°C. All buffers used throughout the process were filtered using membrane filters of 0.22 μm (Millipore). The working concentration for WtIHF and ScIHF was kept as 500 nM in all smFRET experiments.

### Design of DNA Constructs

All the DNA templates used in the present study (labelled or unlabelled) were customised, HPLC purified and purchased from Integrated DNA technologies Inc. (IDT), Coralville, USA. These oligonucleotides were stored at −20°C without any further purification till their use for experiments.

To assemble the two respective Holliday Junctions (H’-HJ and H1-HJ) used for single molecule study, the following component oligonucleotide sequences (34 nucleotides each, except for the biotinylated strand which is 39 nucleotides in length) were used: where the sequences with the same font style correspond to a pair of complementary oligonucleotides. The two Holliday junction constructs used were designed based on the two specific binding sites (H’ & H1) of Wild-type (Wt) and Single-chain (Sc) variants of Integration Host Factor (IHF). So, the two junction constructs differ in terms of the specific protein binding sequence on their H’HJ1 and H1-HJ1 strands respectively. The protein binding sequence in case of each construct has been highlighted in red. The resulting two junctions formed comprised four helical arms of 17 base pairs (bp). Each immobile or non-migratable (incapable of undergoing branch migration due to absence of homologous sequence) was constructed by hybridising four single stranded complementary oligonucleotides. H’-HJ comprises of H’-HJ1, H’-HJCy3, H’-HJCy5 and H-HJ Biotin whereas H1-HJ consists of H1-HJ1, H1-HJCy3, H1-HJCy5 and H-HJ Biotin. The biotinylated strand is common for both constructs. For annealing, the four strands were mixed in a ratio of 1:1:1:1 (Cy3 labelled:Cy5 labelled: Biotin labelled: Unlabelled) to a final concentration of 20 μM in the following Reaction buffers under various experimental conditions as and when stated:

T50 Buffer-10mM Tris, pH 8.0, 50mM NaCl

Buffer 1-10mM Tris, pH 8.0, 50mM NaCl, 2mM MgCl_2_

Buffer 2-10mM Tris, pH 8.0, 50mM NaCl, 50mM MgCl_2_

After mixing the four strands in appropriate buffer as per experimental requirement, annealing was done by heating the sample mixture up to 95°C followed by gradual and controlled cooling down to 8°C (peqSTAR Thermocycler, PEQLAB Biotechnolgie) for 2.5 hours. In order to achieve pico-molar concentration of DNA samples (HJs) for single molecule experiments serial dilutions of the final annealed product in Reaction Buffer according to different experimental conditions (T50/Buffer1/Buffer2). The remaining annealed product was stored in −20°C for future experimental use (if required). For CD and MST experiments, the same unlabelled (without Cy3/Cy5) oligonucleotides were used (Supplementary Table 1)

### Circular Dichroism

CD experiments were conducted using the Chirascan Spectropolarimeter with Spectra Manager version 1.53.00. Raw data was acquired from the instrument and subsequently smoothed using Origin with the Stineman function and a window size of 15. The sample employed for the experiments consisted of H1-HJ at a concentration of 5 µM, dissolved in a 10 mM Tris-HCl buffer with 50 mM NaCl at pH 8.0. The sample solution was loaded into a 1 cm cuvette for measurement. To investigate the effect of MgCl2 on the H1-HJ structure, a titration of MgCl2 was performed by sequentially adding concentrations of 2, 8, 12, 16, and 50 mM into the cuvette. Each addition was followed by thorough mixing, and a new CD spectrum was recorded. The spectra were recorded within the range of 350 nm to 210 nm and averaged over 3 scans to enhance the signal quality. The same experimental procedure was repeated for H’-HJ at a concentration of 5 µM to compare its CD spectra under the same conditions. To examine the binding of proteins and their impact on the junction conformation, WtIHF and ScIHF proteins were added at a final concentration of 5 μM, and CD scans were taken. These experiments were conducted in triplicate to ensure consistency and reliability of the results.

### Quantitative Interaction Studies by Microscale Thermophoresis

Microscale thermophoresis was employed to analyse DNA-protein interactions in this study (Sankhe *et al*, 2023; Joseph *et al*, 2022; Wienken *et al*, 2010; Sparks *et al*, 2022; Gontier *et al*, 2021; Li *et al*, 2022). The proteins WtIHF and ScIHF were purified and labelled using the previously described method. The experiments were conducted using a Monolith NT.115 Blue/Green instrument (NanoTemper Technologies) in three independent replicates. Standard glass capillaries (NanoTemper Technologies) were utilized for the measurements. For the binding studies of WtIHF and ScIHF to both H’-HJ and H1-HJ, the proteins were used at different concentrations in matching protein buffers. The labelled proteins were used at a concentration of 25 nM, while the DNA junction concentrations ranged from 0.98 nM (lowest) to 5 μM (highest). The measurements were performed at 10% MST power. To establish the binding curves, the difference in thermophoresis between the fluorescent molecules in the unbound and bound states was plotted against the ligand concentration. The binding constants (K_d_) were derived from these binding curves. The graphs were represented as fraction bound against ligand concentration. The data presented in the graphs represent three independent experiments and were fitted to a K_d_ binding model, assuming a 1:1 binding stoichiometry.

The proteins WtIHF and ScIHF were labelled with the Lysine dye using The Protein Labelling Kit RED-NHS 2nd Generation. Specifically, 25 μl of DMSO was added to 10 μl of the dye solution before use, and the final concentration of the dye was 600 μM. Subsequently, 50 μl of the 15 μM dye solution was added to 50 μl of the 5 μM protein sample in a microcentrifuge tube. The mixture was carefully mixed by pipetting up and down several times. This resulted in a 100 μl dye-protein solution with an approximate 3-fold excess of dye, which was then incubated for 30 minutes at room temperature in the dark. To purify the labelled proteins, the B-column was filled with assay or equilibration buffer (T50), allowing it to enter the packed resin bed completely by gravity flow. The flow-through was collected and discarded, and this step was repeated three more times. After the incubation of the labelling reaction, 100 μl of the dye-protein solution was directly transferred to the centre of the resin bed in the B-column. Then, 200 μl of T50 buffer was added and allowed to enter the resin bed completely, and the flow-through was collected. Additionally, another 450 μl of T50 buffer was added onto the column, and the flow-through containing the labelled proteins WtIHF and ScIHF was collected by placing a fresh microcentrifuge tube under the column. Finally, the proteins were flash frozen and stored at −80°C until further use.

### SmFRET experiments and Data Acquisition

All single molecule FRET experiments were performed at room temperature (25°C) using our home-built Prism-type Total Internal Reflection Microscopy (p-TIRFM) setup (Figure 1). Pre-drilled quartz slides were taken and chemically processed to make them suitable for DNA immobilization, the details of which is already mentioned in our previous study.

For surface immobilization of DNA samples, Streptavidin at a working concentration of 0.2mg/ml was flown into the narrow microfluidic channels made by sandwiching two thin double sided tape strips between a glass coverslip and PEGylated quartz slide to adhere the two surfaces together. Then, HJs carrying a biotinylated DNA strand were flown into the microfluidic channel and allowed to incubate for 5 mins. After this, the channel was flushed with an appropriate Reaction buffer (as per experimental specification) to wash off any unbound and excess streptavidin. This is followed by introduction of biotinylated samples (diluted to 20pM) into the channel surface coated with streptavidin. After allowing 2-3 minutes of incubation the number of immobilized DNA molecules (∼400) were checked with a single short laser pulse (532 nm). Just before data acquisition (imaging), the channel was flushed with Imaging Buffer comprising of:

i. Reaction Buffer (as per experimental specification)
ii. 5mM Trolox (9-hydroxy-2,5,7,8-tetrmethylchroman-2-carboxylic acid) which quenches triplet state and prevents photo blinking.
iii. Glucose (1μg/ml)
iv. Glucose oxidase (1mg/ml) and catalase (0.03mg/ml) that acts as oxygen scavenger and enhances photostability of fluorophores.
v. Protein (WtIHF/ScIHF)-as per experimental requirement. Protein concentration was kept as 500nM (if and when used).

An inverted fluorescent microscope (Olympus IX71) was used to achieve proper focus over samples. A solid-state diode green (532 nm) laser (Laser Quantum, Stockport, U.K.) was used to excite Cy3 (donor fluorophore) which then transfers its energy to Cy5 (acceptor fluorophore), thereby exciting Cy5 through Forster Resonance Energy Transfer (FRET). The emitted fluorescence signals from Cy3 and Cy5 were then collected simultaneously with the help of water immersion 60X, 1.2 NA, objective (Olympus). Emission signals collected from the fluorophore Cy3 and Cy5 were passed through a 550 nm long-pass filter (Chroma) and eventually separated using 640 nm DCXR dichroic mirror (Chroma). Finally, donor and acceptor emissions were projected onto one half each of the electron-multiplied charge-coupled-device camera (EMCCD, Ixon3+897, Andor Technologies, South Windsor, CT). Movies for all experiments were recorded keeping a multiplication gain of 200 and a frame integration time of 35 milli-seconds (ms) by employing a Visual C++ based data acquisition software.

### Data Analysis

Data from raw movie files were processed using codes written on Interactive Data Language (IDL), a generous gift from Prof. Tae-Hee Lee (Pennsylvania State University, State College, PA). Approximately 200 single molecule traces exhibiting proper anti-correlated intensity between donor and acceptor, with single-step photobleaching and devoid of any photo-blinking events were chosen for next levels of data analysis. Data analysis for these 200 traces were carried out through custom-based codes written on IDL and ensemble histograms were generated. The kinetic rate of transition between various states of all selected molecules exhibiting more than one conformational state was calculated by dividing the output of the transition matrix that was generated from Hidden Markov Model (HMM) code written on IDL by the camera’s exposure time (35 milli-seconds).

The dwell time of such molecules in a particular state was also determined. Apart from ensemble histograms indicating the total number of counts within a particular E_FRET_ range, individual histogram (superimposed FRET efficiency histogram of each of ∼200 individual molecule) was generated with the help of codes written in MATLAB. Gaussian fitting of histogram peaks were done in Origin 2018.

## Acknowledgments

This work is supported by the Department of Atomic Energy (DAE), the Government of India and the Science and Engineering Research Board (SERB-CRG Project no: CRG/2019/006384), India. FI thanks the Council of Scientific and Industrial Research (CSIR), New Delhi, India for providing the scholarship. The authors thank Manali Basu for the scientific discussion and technical help during Circular dichroism experiments.

## Author Contributions

PPM and FI conceptualized the project and designed the work, FI executed the experiments that includes purification of the protein, optimization of the instrument, data acquisition, and data analysis. PPM and DP have interpreted the data. FI has drafted the manuscript and PPM has corrected it.

